# Bed and breakfast in the bush: Selection of resting sites and kill sites by leopards (Panthera pardus) on Namibian farmland

**DOI:** 10.64898/2026.03.18.712594

**Authors:** Nik Šabeder, Teresa Oliveira, Ruben Portas, Lan Hočevar, Urša Fležar, Bettina Wachter, Joerg Melzheimer, Miha Krofel

## Abstract

Sleeping and feeding are crucial for survival of any animal. In case of large predators, knowing where these activities occur can help us understand their behavioural adaptations for coexisting with people and could help mitigating human-carnivore conflicts. Leopard (*Panthera pardus*) is an elusive and highly adaptable large felid that mostly lives outside protected areas and can survive also in close proximity to humans. However, most leopard research in Africa has been conducted in protected areas and we poorly understand leopards’ habitat selection while resting and hunting. To shed light on their coexistence with humans, we investigated habitat features influencing leopard selection of resting and kill sites on farmlands in central Namibia, using generalized linear mixed models (GLMM) under a use-availability study design and blinded field-sampling. Leopards primarily selected resting sites that were located in mountainous, steep, rugged terrain and sites with good concealment while kill sites were selected in mountainous habitats. Human infrastructure did not affect leopard resting and kill site selection. Thus, the capacity of leopards to perform essential life-supporting behaviours while coexisting with people appears to be primarily driven by their ability to remain concealed, rather than spatially avoiding humans.

## INTRODUCTION

Resting and foraging are among the essential animal behaviours with direct fitness implications (Kusler et al. 2017). However, these are also behaviours during which animals are most vulnerable to human disturbance or predation by larger competitors (Gorman et al. 1998; Kusler et al. 2017; Hočevar et al. 2021). Resting and foraging are particularly relevant for large carnivore species, which are disproportionately affected by human persecution, due to low population densities and apex trophic position (Ripple et al. 2014; Darimont et al. 2015; Krofel et al. 2015). Accordingly, identifying micro-scale habitat requirements of particular behaviours can play a vital role in ensuring adequate land-use management within large carnivore home ranges (Kolowski & Woolf 2002).

To avoid interactions with humans, many large carnivores have adopted nocturnal lifestyles, particularly in areas with frequent daytime human activity (Burton et al. 2024). Therefore, carnivore species seek resting sites that offer safety from people and other potential threats, as well as shelter from environmental factors such as rain, sun, and wind (Wam et al. 2012). Consequently, resting sites are often selected in concealed, inaccessible locations in elevated and rugged terrain with a presence of special relief features and a good overview on their surroundings (Kusler et al. 2017; Skuban et al. 2018; Hočevar et al. 2021; Čonč et al. 2024) and located at sites that provide optimal micro-climate conditions (Jackson 1996).

Large carnivores select foraging areas according to two tactics. They either focus on hunting where it is easier to capture prey (landscape hypothesis) or in areas where prey is more abundant (prey abundance hypothesis) (Balme et. al 2007). Both tactics are supported in the literature; for example, lions (*Panthera leo*) preferred areas with dense vegetation that provides stalking cover (Hopcraft et al. 2005), leopards (*Panthera pardus*) favoured intermediate degree of vegetation (Balme et al. 2007), whereas grey wolves (*Canis lupus*) favoured more open habitats, where it was easier to detect prey (Hebblewhite et al. 2005). On the other hand, Iberian lynx (*Lynx pardinus*) and mountain lions (*Puma concolor*) focused their hunting on areas with highest prey abundance (Pike et al. 1999; Palomares et al. 2001), whereas Eurasian lynx (*Lynx lynx*) appear to hunt regardless of relative prey abundance (Nilsen et al. 2009; Krofel et al. 2014). Since large carnivores spend substantial time and energy in hunting, a secure access to their kills and concealment from scavengers and people is needed to justify this investment, particularly when feeding time is prolonged, as is the case with many solitary predators (Krofel et al. 2008, 2012; Balme et al. 2017). This underscores the importance of selecting specific landscape features to make and consume a kill (Mukherjee and Heithaus, 2013; Tallian et al. 2023).

Many large carnivores are specialised in hunting ungulates, which in human-dominated landscapes often brings them into conflict with people due to the risks associated with depredation of domestic livestock, as well as valuable wildlife species (Treves and Karanth 2003). Knowing where predators preferably kill their prey can help adjusting livestock husbandry practices to reduce the risk of human-carnivore conflicts (Treves et al. 2011; Melzheimer et al. 2020). Thus, studying the selection of kill sites by large carnivores can be important not only for understanding their ecological requirements, but also for their abilities to coexist with people, which can facilitate the development of effective conservation strategies. However, for many threatened species of large carnivores such knowledge remains limited.

Leopards are large, solitary felids occurring throughout most of sub-Saharan Africa and southern Asia (Bailey 1993; Rodríguez-Recio et al. 2022). They are highly adaptable predators, able to survive in a range of habitats from deserts to savannahs and rainforests, as well as capable of living in close proximity to humans and human infrastructure, sometimes even on the outskirts of large cities (Athreya et al. 2013; Gubbi et al. 2020). However, in the recent decades many leopard populations have declined due to habitat loss, fragmentation, and human-leopard conflict (Jacobson et al. 2016). The species currently occupies only 25–37% of its historic range (Jacobson et al. 2016) and is listed as vulnerable according to the International Union for Conservation of Nature (IUCN, Stein et. al. 2020). The vast majority (87%) of the remaining distribution range is located outside of protected areas (Stein et al. 2011; Jacobson et al. 2016). Generally, leopard activity and habitat use are affected by an interplay of anthropogenic disturbance, presence and activity of dominant competitors and prey (Mugerwa et al. 2017; Allen et al. 2020). However, most of the previous research focusing on leopard ecology has been limited to protected areas (Balme et al. 2014), underscoring the knowledge gap in large part of the species’ distribution range, where also conflicts with humans and potential persecution is highest. Balme et al. (2007) reported that leopards hunting in a protected area in South Africa favoured habitats with intermediate vegetation cover, while prey abundance appeared less important. But beyond these factors, little is known about habitat characteristics influencing the distribution of leopard kills and to our knowledge no studies were conducted on the selection of day resting sites by leopards.

In this study, we aimed to explore selection of resting sites and kill sites by leopards in the Namibian farmlands, where this predator coexists with people, livestock (predominantly cattle) and wild prey. Specifically, we investigated the influence of several environmental factors and human infrastructure on leopard selection of resting and kill sites using a use-availability study design. Based on previous research on leopards and other large felids (e.g. Jackson 1996; Balme et al. 2007; Krofel et al. 2008; Kusler et al. 2017; Hočevar et al. 2021; Čonč et al. 2024), we predicted that leopards select resting sites in inaccessible, rugged and mountainous terrain with steep slope, presence of special relief features, a good concealment and at the same time good (elevated) overview of their surroundings. We predicted that leopards preferentially kill large prey in intermediate vegetation cover with good concealment from scavengers and people and on locations with steeper slope, on rugged and mountainous terrain. We also predicted that leopards in the farmland would avoid human infrastructure when selecting a resting site or kill site, given that human persecution is the main mortality factor in this landscape (Swanepoel et al., 2015).

## MATERIAL AND METHODS

### Study area

The study was conducted on freehold farms in central Namibia (23°01’-21°26’ S and 16°51’-18°17’ E). The region consists of large flat plateaus interspaced with hills and rugged mountains in the Khomas Highlands and Auas Mountains, formed by an undulating to steep hilly and mountainous landscape, generally with shallow, stony soils. The study area includes dry riverbeds, high mountains, rocky hills and lowlands. Elevation ranges between 1,680 and 2,479 m a.s.l. Annual precipitation ranges from 300 to 350 mm per year, with rain concentrated from approximately November to May. Maximal day temperatures are up to 35°C in the rain season and can get as low as frost in the night in cold dry season, with an average annual temperature of 16 - 18°C (Mendelsohn et al. 2022). The vegetation in the study area consists of the highland savannah, dominated by *Acacia hereroensis* and over 50 grass species. The woody vegetation also contains *Acacia mellifera, A. karroo*, *Ziziphus mucronata, Combretum apiculatum, Tarchonanthus camphoratus* and *Catophractes alexandri* (Mannheimer and Curtis 2010; Strohbach, 2017). Eland (*Taurotragus oryx*), greater kudu (*Tragelaphus strepsiceros*), mountain zebra (*Equus zebra*), red hartebeest (*Alcelaphus buselaphus*), gemsbok (*Oryx gazella*), warthog (*Phacochoerus africanus*), springbok (*Antidorcas marsupialis*) and steenbok (*Raphicerus campestris*) are the most common ungulates and potential leopard prey species (Voigt et al., 2018; Melzheimer et al. 2020). Beside leopards, cheetahs (*Acinonyx jubatus*) and brown hyenas (*Parahyaena brunnea*) also inhabit the area. The main land use is cattle and game farming, horse breeding, trophy hunting and tourism. Namibia is home to one of the largest populations of leopards in the world, inhabiting most of the country except for the highly populated northern region, the arid south-east farmlands and the desert coast (Richmond-Coggan, 2022). Most of the leopards in the country live outside of protected areas and our study area is assessed to contain one of the highest leopard densities in Namibia (Richmond-Coggan, 2022).

### Leopard capture and chemical immobilization

Leopards were captured between 2012 and 2022 throughout the year using mechanical or electronical metal box traps (Portas et al. 2019). The traps were set at known scent-marking locations or at other frequently-used locations, identified by camera-trapping and searching for leopard signs. Target individuals were attracted to the capture sites with meat and/or audio baits with sounds of prey species or leopard vocalizations. Leopards were immobilized with a dart gun using 0.04-0.06 mg/kg medetomidine hydrochloride (Medetomidine 10 mg/mL; Novartis, Johannesburg, South Africa) and 2.5-3.0 mg/kg ketamine (Ketamine 1G; Kyron Laboratories, Johannesburg, South Africa) and reversed with 2.0-2.4 mg/kg atipamezole (Antisedan®; Novartis). Leopards were fitted with GPS remote-download (e-obs GmbH, Germany) or satellite collars (Vectronic Aerospace GmbH, Germany, or Followit Wildlife, Lindesberg AB, Sweden) with three-axes accelerometers. The collars always weighed less than 2% of the body mass of the leopards and were set to attempt to obtain 6-48 GPS positions per day.

### Identifying leopard resting sites and kill sites

We used GPS data from 28 collared leopards to locate leopard resting and kill sites (Online Resource Material 1, Table S1). Resting sites were defined as daytime GPS fixes at midday (12:00 local time), when leopards were inactive, which was verified with information from accelerometers sensors. Specifically, activity levels of 0 – 11 % of the maximum activity around the midday position (+/- 10 min) were considered resting behaviour (Heurich et al. 2014; Hočevar et al. 2021). Kill sites were identified using GPS location cluster (GLC) analysis (Merrill et al. 2010). Kill sites were generally selected for verification in the field when clusters of GPS points formed within a 300 m radius from the center of the cluster spanned at least 24 hours, but occasionally we also visited shorter clusters (8.8% of all clusters checked). Sometimes (5.6%, n=373 kills) we observed that leopards dragged prey remains from the site of the kill to feeding sites for 2-120 m (mean = 28 m). In these cases, we sampled the location where the killing was assumed to take place. When no drag marks were observed, we assumed that the killing took place at the location where prey remains were found. In our study area leopards rarely (2.6 %) hoisted their prey on trees.

### Field sampling, data processing and covariate selection

To investigate the selection of resting and kill sites by leopards, we applied a use-availability study design at the home range level (i.e. 3^rd^ order of selection; Johnson 1980). This approach compares two samples: known (used) locations and random (available) locations. We considered the resting sites and kill sites as “used” sample and random locations as “available” sample. We visited resting sites, kill sites and random locations to extract the covariates in the field, that could not be extracted remotely (see below) in accessible areas, and not equally across the whole home ranges, because of limited accessibility to some of the private farms. We estimated the home range size as 95% minimum convex polygon (MCP).

To reduce potential bias during the extraction of covariates in the field, all field researchers were unaware (blinded) during the fieldwork to whether a site we were sampling was a leopard resting sites or a random location.

We considered 14 covariates potentially influencing leopard resting and kill site selection, divided in the four groups: topography, visibility, vegetation, and human infrastructure (Online Resource Material 1, Table S2). Within topography, we considered relief type, special relief features, terrain ruggedness, and orientation as categorical covariates, and slope as continuous covariates (Online Resource 1, Table S2). Relief type included “flat”, “hillside” and “mountainous” habitats. Special relief features described the presence of “hillside”, “hilltop”, “ridge”, “small cliff”, “large cliff”, “cave”, or “none” within 30 m radius from the location. Terrain ruggedness described ruggedness within 20 m radius from the location, as “flat”, “medium rugged”, and “very rugged”. Orientation direction indicated the direction of the slope the visited site was facing, and included south, southeast, southwest, east, west, north, northeast, and northwest, or we labelled it as flat, when site was not located on a slope.

We considered three covariates related to the visibility at the site: concealment (continuous), long-distance view (continuous) and long-distance viewpoint (categorical). We measured the concealment of each site following Hočevar et al. (2021) by placing a backpack on a site, and walking in the four cardinal directions, counting the steps (0.75 m in length) until we could no longer see the backpack. We then averaged values for all of the directions. Long-distance view described the amount of an unobstructed view (in degrees) of the landscape between up to 1000m from the site. Long-distance viewpoint described the viewpoint of an animal/the observer from the site. It included “elevated”, “flat”, “looking up from below” and “none” (Online Resource Material 1, Table S2).

For vegetation cover, we included “habitat openness” and “canopy cover” (both categorical). Habitat openness included “open” (0-25 % of vegetation cover), “semi open” (26-50 %), and “dense” (51-100 %) and described vegetation cover with bushes and trees. Canopy cover described vertical cover of all vegetation directly above the site: “open” (0 % vegetation cover), “little open” (1-25%), “semi open” (26-50%) and “dense” (> 50 %). Additionally, we considered the availability of midday shade if there was shaded ground available at least in the size of a leopard body size provided by a large tree, bush or cliff.

We considered three covariates for human infrastructure. We calculated the Euclidean distance from the visited site to the nearest main public road, waterhole or other artificial water body with water present for the majority of the year, and inhabited house. We calculated these three distances from a vector shapefile including roads, waterholes and houses, which we mapped with satellite imagery. We used the “Euclidean distances” tool from QGIS Desktop 3.16.5 (QGIS Development Team, 2023).

For the animals collared from 2021-2022 (Online Resource Material 1, Table S1) we extracted the parameters of the resting sites in the field (i.e., presence of special relief features, orientation, concealment, long-distance view, long-distance viewpoint, habitat openness, canopy cover; Online Resource Material 1, Table S2). Same parameters were extracted at kill sites between 2014 and 2022 from 16 leopards. For the remainder of the data, we assessed parameters remotely (i.e., relief type, slope). Therefore, we conducted our analyses with two datasets: resting and kill sites from all leopards (n=28), and resting and kill sites for leopards (n=5 for resting sites; n=16 for kill sites) with data extracted in the field (Tables 1; Online Resource Material 1, S1).

For the analyses with all 28 collared individuals, we considered only the two explanatory covariates that remain constant throughout the years and are available from the remote-sensing datasets, i.e. relief type and slope. We included 4,102 leopard resting sites identified with GPS data as “used” sample and selected 41,020 random locations within the 95% MCP leopard home ranges as “available”, i.e. 10-times the “used” sample (Northrup et al. 2013). We followed the same procedure for the 373 identified kill sites (Table 1). When using data extracted in the field from five leopards for resting sites and 16 leopards for kill sites, we included in addition to relief type and slope also 14 further explanatory covariates which we extracted in the field (Online Resource Material 1, Table S2). These 14 additional covariates better described the microhabitat characteristics, but could change between the years, therefore we did not collect such data for sites used by the leopards from more than a year prior to extraction in the field. In total, we visited 192 leopard resting sites, 373 kill sites, and 200 random locations sampled within the 95% MCP leopard home ranges (n=28).

### Modelling approach

We built generalised linear mixed models (GLMMs) with a binary response variable representing “use” and “available” locations (1 and 0, respectively). We included leopard individual ID as a random effect in all models to account for the unbalanced sample sizes (Gillies et al. 2006). We built different models using different combinations of fixed effects to investigate various hypotheses (Table 1). We ranked models according to the Akaike’s Information Criteria (AIC) and considered models with ΔAIC lower than 2 from the lowest AIC value as the best models (Burnham and Anderson 2002). We also added a null model to evaluate the gain of adding a particular set of covariates. Before building the models, we checked for correlation between each pair of covariates by using the variance inflation factor (VIF), and we excluded covariates from the model if the VIF was > 3 (Zuur et al. 2010). We assessed model fit following the method proposed by Boyce et al. (2002), i.e. by extracting the model parameters and plugging them into a RSF (resource selection function), and then employed *k*-fold cross-validation. Specifically, for the latter, we partitioned the data, using 80% for model training which were then used for predicting the probability of use for the remaining 20% of data. The process was repeated five times, ensuring all data were used. To evaluate the predictive performance, we calculated Spearman rank correlations between the frequency of cross-validated used locations and ten probability bins of similar size, which represented the range of predicted values. According to Boyce et al. (2002), a model with strong predictive power should exhibit a high correlation (ρ > 0.80).

The GLMMs built with the subsampled dataset of kill sites (i.e., with field-extracted data) showed that the considered covariates did not influence selection of kill sites by leopards (i.e. the null model had the lowest AIC value; Table 2). However, we still performed bivariate analyses to obtain preliminary information regarding a potential effect of these covariates on kill site selection. We used Student’s *t*-test for continuous covariates and Pearson χ^2^ statistics to test the significance of categorical covariates. For these analyses, we used a subset of the field-extracted kill sites (n=195) for which we had data on the covariates extracted in the field (i.e., presence of special relief features, orientation, concealment, long-distance view, long-distance viewpoint, habitat openness, canopy cover; Online Resource Material 1, Table S2). We compared these with similarly sampled random locations (n=195) within the same spatial range of the kill sites distribution.

All statistical analyses and data visualisation were conducted using software R version 4.1.3 (R Core Team, 2023), packages ggplot2 v3.5.1 (Wickham 2016), lme4 v1.1-21 (Bates et al. 2015), forcats v1.0.0 (Wickham 2016) and dplyr v1.1.4 (Wickham et al. 2023).

**Table 1.**
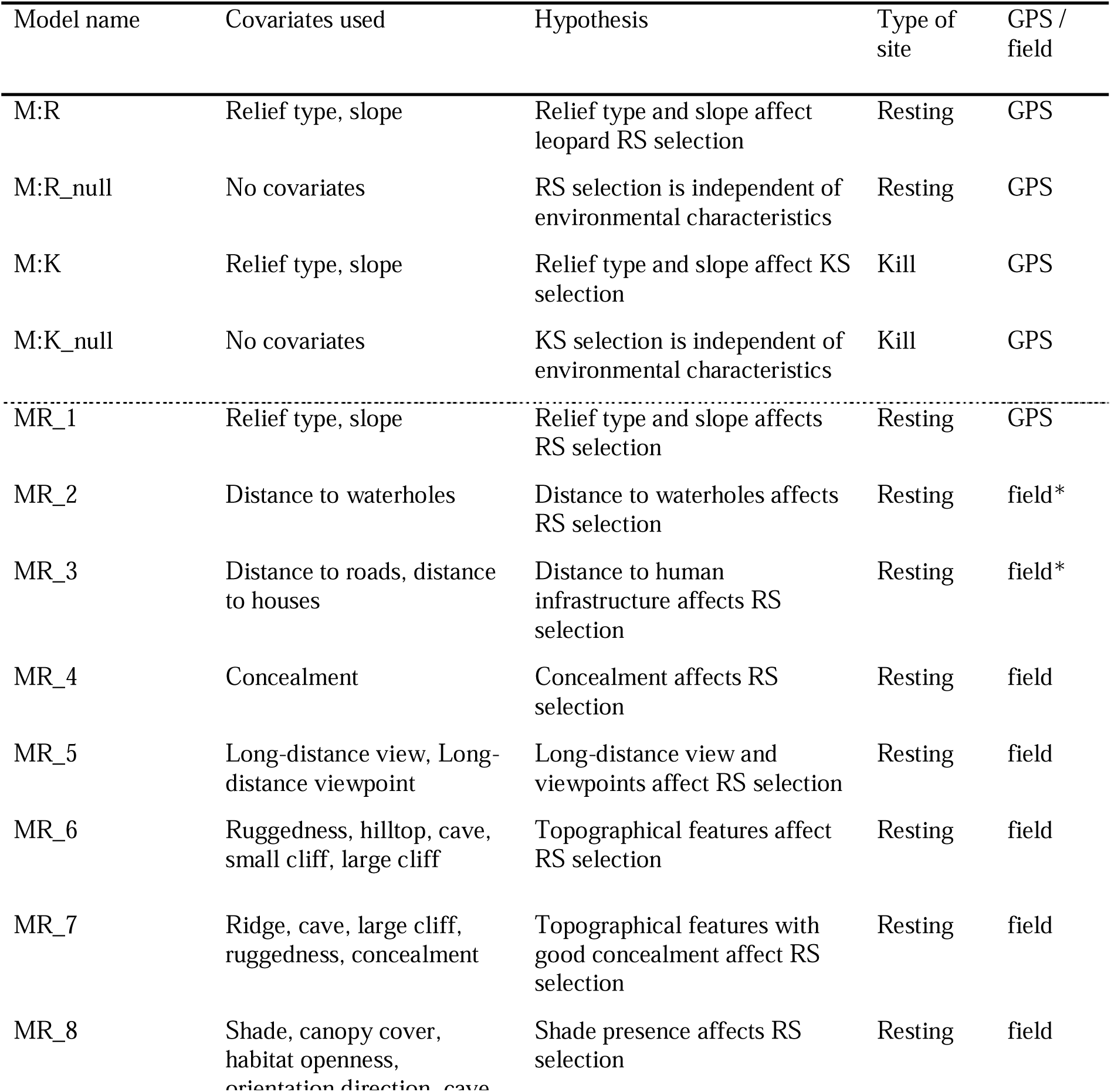
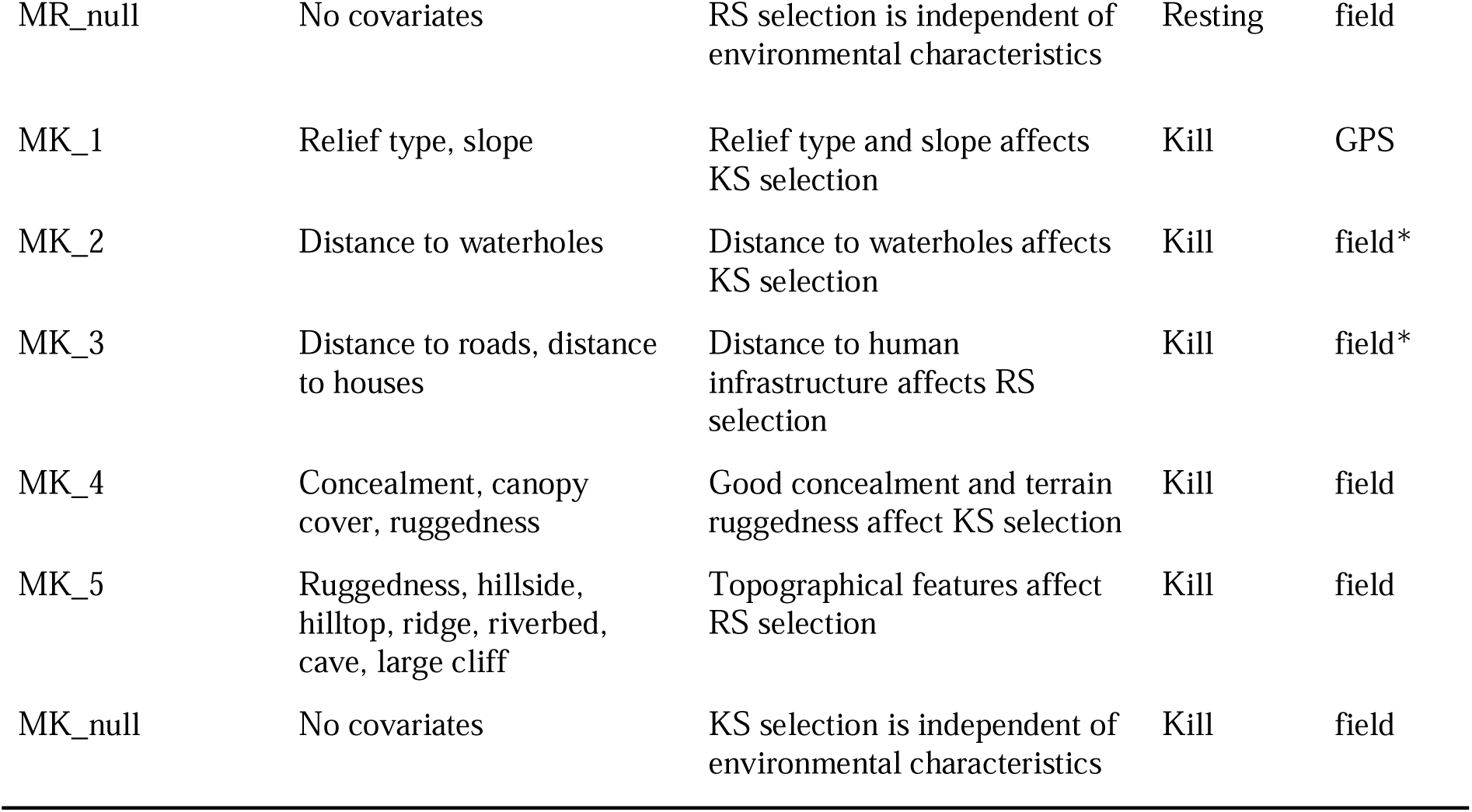
Generalized linear mixed models (GLMMs) built to describe leopard resting site (RS) and kill site (KS) selection. Dashed line divides the models in two parts: (1) four models for the selection of resting sites and kill sites for which GPS data from 28 leopards and the covariates relief type and slope based on remote sensing data were used; (2) nine models for the selection of resting sites for which GPS data from five leopards for resting sites and 16 leopards for kill sites and field extracted measurements on 14 covariates were used. In the last column, “GPS” refers to remotely-measured locations that we did not visit in the field, “field” refers to locations we visited and measured the parameters in the field and “field*” represents locations we visited and afterwards measured distances remotely with GIS.

## RESULTS

### Resting sites

The model that best described leopard (n=28) selection of resting sites (n=4,102) included relief type and slope (Table 2). Specifically, leopards selected steeper slopes and hillsides and mountains as opposed to flat areas (Table 3). When considering also microhabitat characteristics extracted in the field, the model that best described the selection of the field-extracted resting sites (n_sites_=192, n_individuals_=5) included special relief features (ridge, cave, large cliff), terrain ruggedness and concealment (Table 2). The coefficients of the best model (Fig 1) suggested that leopards selected rugged terrain with good concealment for their resting sites.

**Table 2:** GLMMs for leopard resting site (RS) and kill site (KS; separated by thin line) selection in central Namibia. Dataset column represents which dataset was used. “All” means the entire dataset with remotely assessed resting sites or kill sites (representing the “use”), and random locations (representing the “availability”) of 28 leopards with slope and relief type as covariates. “Field-extracted” represents dataset with resting (for 5 leopards) or kill sites (for 16 leopards) and random locations (representing “use” and “availability”, respectively) with 14 covariates. Model names (see Table 1 for details), *k*-number of covariates in the model, AIC (Akaike Information Criterion), ΔAIC (difference of AIC scores of each model in respect to the best model) and ρ—Spearman rank correlation values are presented. Best models are highlighted in bold. Coefficients for M:R, M7 and M:K are given in Fig 1 and Tables 3 and 4.

| Dataset | KS or RS | Model | $k$ | AIC | ΔAIC | $\rho$ |
| --- | --- | --- | --- | --- | --- | --- |
| **All** | **RS** | **M:R** | **4** | **394.67** | **0** | **0.84** |
|  |  | M:R_null | 2 | 409.42 | 14.75 | 0.81 |
| **Field-extracted** | **RS** | **MR_7** | **8** | **456.84** | **0** | **0.76** |
|  |  | MR_4 | 3 | 467.94 | 11.1 | 0.71 |
|  |  | MR_8 | 14 | 481.53 | 24.69 | 0.64 |
|  |  | MR_6 | 8 | 516.98 | 60.14 | 0.81 |
|  |  | MR_1 | 5 | 529.51 | 72.67 | 0.69 |
|  |  | MR_2 | 3 | 541.59 | 84.74 | 0.79 |
|  |  | MR_null | 2 | 547.29 | 90.45 | 0.54 |
|  |  | MR_5 | 3 | 548.99 | 92.15 | 0.63 |
|  |  | MR_3 | 4 | 550.77 | 93.93 | 0.82 |
| **All** | **KS** | **M:K** | **4** | **387.72** | **0** | **0.86** |
|  |  | M:K_null | 2 | 411.38 | 23.66 | 0.83 |
| **Field extracted** | **KS** | **MK_null** | **2** | **487.23** | **0** | **0.68** |
|  |  | MK_4 | 5 | 511.72 | 24.49 | 0.63 |
|  |  | MK_2 | 3 | 519.65 | 32.42 | 0.70 |
|  |  | MK_1 | 4 | 527.39 | 40.16 | 0.69 |
|  |  | MK_5 | 8 | 533.47 | 66.24 | 0.72 |
|  |  | MK_3 | 4 | 562.86 | 75.63 | 0.61 |

**Table 3:** GLMM coefficients for the best model (M:R in Table 2) describing the selection of resting sites (“used” sample) by leopards in comparison with random locations (“available” sample). Reference for the relief type was “flat”.

| Model | Covariate | Estimate | Std. err | *p* |
| --- | --- | --- | --- | --- |
| M:R | Intercept | -2.590 | 0.29520 | <0.001 |
|  | Slope | 1.429 | 0.167 | <0.001 |
|  | Relief_Hillside | 0.513 | 0.443 | <0.001 |
|  | Relief_Mountain | 1.063 | 0.812 | <0.001 |

**Fig 1:**
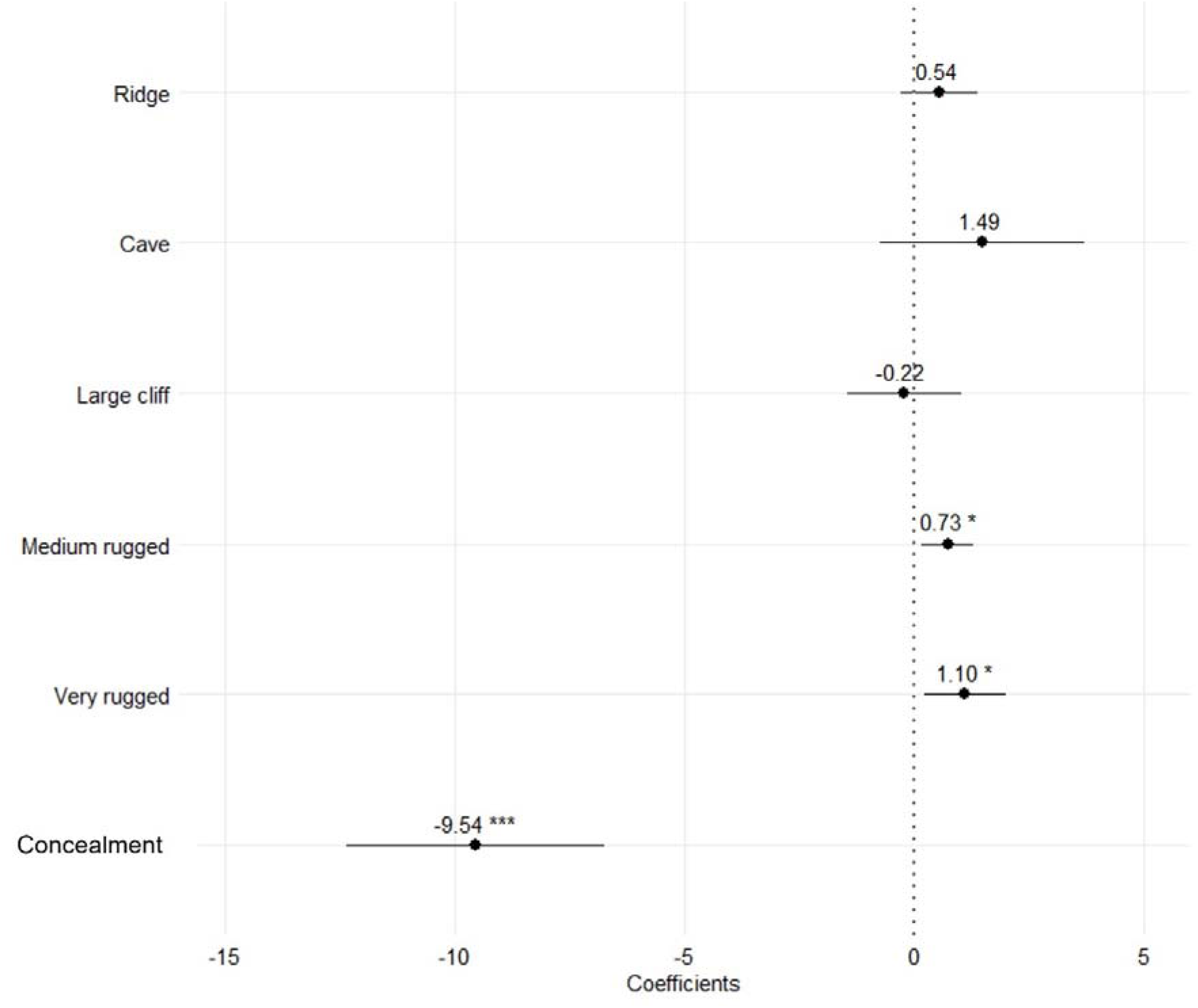
Coefficients from the best model (MR_7 in Table 2), describing leopard resting site selection in respect to availability. Ridge, Cave and Large cliff are binary categories of the covariate “Relief type”, while Medium and Very rugged are categorical covariates, with “flat” as reference category of the covariate “Terrain ruggedness”, and Concealment is a continuous covariate. Model coefficients and 95% confidence intervals for each covariate are provided, with significant covariates (i.e. intervals not containing zero) marked with * symbols.

### Kill sites

The model that best explained the selection of the leopard kill sites (n=373) when considering the whole dataset (28 individuals) and remotely assessed data, included slope and relief as covariates (Table 2). Specifically, leopards killed their prey significantly more often on hillsides and mountains compared to flat areas when tested against random locations as availability sample (Table 4). Slope did not significantly influence the location of kill sites (Table 4).

**Table 4:** GLMM coefficients for the best model (M:K in Table 2) describing the selection of kill sites (“used” sample) by leopards in comparison with random locations (“available” sample). Reference for the relief type was “flat”.

| Model | Covariate | Estimate | Std. err | *p* |
| --- | --- | --- | --- | --- |
| M:K | Intercept | -1.881 | 0.627 | 0.003 |
|  | Slope | 0.292 | 0.468 | 0.533 |
|  | Relief_ Hillside | 1.071 | 0.161 | <0.001 |
|  | Relief_Mountain | 1.417 | 0.288 | <0.001 |

When looking at the results from bivariate analyses, leopards (n=16) preferably killed prey in the mountains, dense vegetation, and rugged terrain compared to random locations (n_killsites_=195; n_randomlocations_=195; Pearson χ^2^ test, p ≤ 0.002; Online Resource Material 2, Fig S1). Kill sites were also found, at locations with more concealment, closer proximity to farmhouses, and greater distance from waterholes compared to random locations (t-test, p ≤ 0.001; Online resource Material 2, Fig S2). Distances to roads was not different between the random sites and kill sites (p = 0.12).

## DISCUSSION

Our findings indicate that leopards on Namibian farmland preferably rest and kill prey at difficult-to-access sites that offer good concealment to avoid detection, rather than to spatially avoid humans. For resting sites, leopards selected steep and mountainous terrain, while for kill sites they selected mountainous terrain. Concealment and ruggedness were also crucial habitat features for leopard resting site selection. This emphasizes the critical role of topography and vegetation in shaping the spatial ecology of leopards, as well as the species’ ability to coexist with people.

Observed preference for resting sites with good concealment aligns with the leopards’ inherent need and ability to remain concealed. Similarly, the bivariate analyses indicate that the same applies also for kill sites. This is specifically relevant when trying to evade potentially dangerous encounters with dominant competitors (Bailey 1993; Rafiq 2016; Ramesh et al. 2017; Searle et al. 2021) or humans (Ngoprasert et al. 2007; Odden et al. 2014). Although our study is the first one focussing on leopard selection of resting and kill sites, results are similar to previous observations on general movement and habitat selection in leopards. For example, leopards in a human-dominated area in Kenya generally selected areas that provided concealment, including during the daylight hours (Van Cleave et al. 2018). Similarly, in India leopards were using high sugar cane fields or patches of scrub, which provided a daytime cover (Odden et al. 2014). Also, a habitat-use analysis by Kshettry et al. (2017) demonstrated that dense vegetation and lower density of houses were the strongest positive predictor of Indian leopard habitat use in a densely-populated landscape.

### Resting sites

Preference for rugged, rocky and steep terrain with good concealment that we observed in leopard resting sites is similar to habitat selection of many other solitary felid species. For example, Eurasian lynx selected resting sites at locations close to rocky outcrops, in steep and rugged terrain and with low visibility (Podgórski et al. 2008; Hočevar et al. 2021; Signer et al. 2019; Čonč et al. 2024). Also, pumas selected sites with dense canopy cover, with dense vegetation cover, and in a rugged habitat (Kusler et al. 2017), snow leopards (*Panthera uncia*) showed a strong preference for rugged, steep, and rocky terrain, often near vegetation or cliffs (Jackson 1996), and European wildcats (*Felis silvestris*) selected vegetation shelter structures near forest edges (Jerosch et al. 2010). In contrast to these solitary felids, adult territorial lions preferentially rested in sites with high visibility and in open habitats (Elliot 2013). This might be connected with the dominant position that group-living lions have in the African carnivore guild, while leopards and other solitary felids require safety from dominant competitors, similar to dispersing lions that need to remain undetected by the territorial conspecifics.

In contrast to our expectation, leopards did not avoid human infrastructure during resting. This differs from the conventional notion that human presence deters leopard habitat use (Martins and Harris 2013; Strampelli et al. 2018, 2022) and might be explained by relatively low human density on Namibian farmland. Similar contrasting results for resting site selection were reported for felids in Europe, where Eurasian lynx avoided human infrastructure during the day (Belotti et al. 2018; Čonč et al. 2024) while the European wildcats did not (Jerosch et al. 2010).

Besides providing concealment, steep and rugged terrain also provides opportunity for good overview of the surroundings, enabling predators to spot potential prey and/or detect approaching humans or other threats (Hočevar et al. 2021). Nevertheless, our prediction that leopards select resting sites that provide good long-distance view was not well supported. This suggests that leopards’ priority is to remain concealed, possibly also from approaching prey, rather than to spot prey or danger from a distance. This might explain the selection of hilltops, ridges and cliffs by other solitary felids (Jackson 1996; Hočevar et al. 2021; Čonč et al. 2024) and selection for open terrain by adult lions (Elliot 2013).

A limitation of our study was that the resting sites visited during the field extraction were limited to the last two years of our study. Therefore, for most covariates we could only perform analyses on five individuals. Nevertheless, we observed similar selection patterns between these five and all 28 collared individuals for the two topographical covariates (slope and relief) available from remote sensing, suggesting that habitat selection of resting sites by our subset leopard dataset could be representative for the larger population in the study area.

### Kill sites

Selection of kill sites showed similar patterns to the selection of resting sites when using the whole dataset and were partially in line with our predictions, as we observed that leopards preferentially killed prey in mountainous and terrain, but we also observed that slope did not play a significant role. Despite not having model support for our subset dataset of field-checked kill sites, we ran basic statistical analyses to get the first impressions and insights into the locations where kills occurred. According to the bivariate analysis, there was indication for selection of concealed, rugged and densely-vegetated terrain. Beside advantages that such habitats provide for hunting, this selection might also reflect avoidance of human persecution, which is widespread in our study area and represent the main mortality cause (unpublished data). Leopards are stalking and ambush predators (Bailey 1993), thus choosing sites that provide good concealment can enhance their chances of remaining hidden and approach prey close enough for a successful capture. This is partly consistent with the findings of Balme et al. (2007) from a protected area in South Africa.

Despite finding significant differences in the proximity of random vs. kill sites to farm houses and waterholes, these features are unlikely to have had an ecological effect due to large mean distances of leopard kill sites to those features (> 2.5 km; Online Resource Material 2, Fig S2). However, if we assume nonetheless that there would be an ecological effect this would again indicate that leopards rely on remaining concealed to avoid detection rather than to avoid areas frequented by people. Additional adaptation of leopards to remain undetected by people in the farmland is to kill and consume prey in a dense bush, rather than to hoist it on a tree, where it would be more easily observed by humans. Lack of spatial avoidance of houses and roads when killing prey might also bring certain foraging advantages to the leopards. For example, risk of kleptoparasitism by larger scavengers can be lower when being closer to human infrastructure (Robins et al. 2025), and anthropogenic noise (e.g. from major roads) may facilitate leopards approaching prey undetected. Several of the kills we found were located in direct proximity to major (B1) road that has loud traffic also in the twilight and nighttime hours.

However, the results that relied on bivariate analysis need to be treated with caution, as confounding factors could have influenced the results. Therefore, further research based on larger sample sizes that would allow for multivariate analysis is recommended. To avoid excessive field effort, this could be achieved also using remote-sensing techniques and high-resolution environmental data, such as LiDAR (Čonč et al. 2022, 2024), when they will become available for more parts of the leopard range.

## Conclusions and implications

Our research on the selection of resting and kill sites is the first study focussing on this essential part of leopard ecology. Our results demonstrate that the driving factor in selecting habitats for resting and, to a lesser extent, foraging, is the need to for concealment and inaccessibility, which can be provided by steep, rugged terrain and/or dense vegetation. When such habitat features enable them to mitigate detection risk, leopards can adapt and persist in proximity of people, as long as they are being tolerated and persecution does not exceed unsustainable levels (Athreya et al. 2013). This elusive behaviour might explain their unique ability among the *Panthera* cats to survive even on the edges of large cities, such as Mumbai in India (Braczkowski et al. 2018).

However, human impact on natural habitats is increasing throughout the world (Yang et al. 2024). In some areas, destruction of habitat features that provide concealment can make it increasingly difficult for species like leopards and other large carnivore species to find a secure site for resting or consuming a prey. This is especially relevant for leopards, for which the majority of distribution range is situated outside of protected areas, where they often interact with humans and livestock (Stein et al. 2011; Jacobson et al. 2016). Thus, targeted conservation measures to protect essential habitat features from excessive human disturbance may be required (Athreya et al. 2013; Čonč et al. 2024). Based on our results, when creating new protected areas to facilitate leopard recovery, it would be advisable to focus on rugged, hilly and mountainous areas, and when such habitats are not available, to focus on habitats with dense vegetation.

Finally, our results can be used to assess the risk for livestock depredation and to design mitigation measures for this widespread conflict. For example, observed preferences for leopards to kill prey in the mountainous terrain suggest that keeping vulnerable domestic animals (e.g., small livestock or calves) in flat and open areas could help reduce farmers’ losses to leopards, where such habitat heterogeneity is available in the landscape. Similar approaches of incorporating novel insights from ecology of large carnivores into livestock husbandry practices have already proved successful in facilitating coexistence between extensive farming and native predators (Melzheimer et al. 2020). Therefore, we recommend further research into behaviour-specific habitat selection, especially for the conflict-prone species.

## Supporting information

Online Resource Material

Online Resource Material 3

## ACKNOWLEDGMENTS

We thank the Ministry of Environment, Forestry and Tourism of Namibia for permission to conduct the study. We are grateful to the Namibian farmers, including members of the Seeis and Auas Oanob Conservancies for cooperation, giving us access to their farms and joining us for fieldwork. We are especially grateful to the Freyer family from farm Claratal and the farm Krumhuk community for extensive help with the fieldwork on their farms and hosting us during the fieldwork. We also thank Dirk Bockmuhl and Vera Menges for help with capturing leopards and downloading telemetry data.

## DATA AVAILABILITY

The datasets generated during the study is included as Online resource material (Online resource material 3).

## COMPLIANCE WITH ETHICAL STANDARDS

### Ethical approval

All relevant international, national, and institutional guidelines for the care and use of animals were adhered to, ensuring compliance with the legal requirements of the host country. All capture and immobilization procedures were approved by the Internal Ethics Committee of the Leibniz Institute for Zoo and Wildlife Research (IZW, Berlin, Germany) (permit number: 2002-04-01) and the Ministry of Environment, Forestry and Tourism of Namibia (permit numbers: 1689/2012, 1813/2013, 1914/2014, 2067/2015, 2194/2016, 2208/2017 and RCIV000822018).

## AUTHOR CONTRIBUTIONS

Conceptualization: Nik Šabeder, Teresa Oliveira, Miha Krofel; Methodology: Nik Šabeder, Teresa Oliveira, Miha Krofel; Formal analysis and investigation: Nik Šabeder, Teresa Oliveira, Ruben Portas, Lan Hočevar, Urša Fležar, Miha Krofel; Writing - original draft preparation: Nik Šabeder, Miha Krofel; Writing - review and editing: Nik Šabeder, Teresa Oliveira, Ruben Portas, Lan Hočevar, Urša Fležar, Bettina Wachter, Joerg Melzheimer, Miha Krofel; Funding acquisition: Bettina Wachter, Joerg Melzheimer, Miha Krofel; Resources: Bettina Wachter, Joerg Melzheimer, Miha Krofel, Supervision: Joerg Melzheimer, Miha Krofel.

## STATEMENTS AND DECLARATIONS

### Funding

The study was funded by the Slovenian Research and Innovation Agency (grants no. J1-50013, N1-0163, P4-0059 and MR-Šabeder), the Messerli foundation in Switzerland, the German Academic Exchange Service and the Leibniz Institute for Zoo and Wildlife Research in Germany.

### Conflict of interest

The authors declare that they have no conflict of interest.

### Disclaimer

All sources of funding are acknowledged and we do not expect any direct financial benefits that could result from this publication.

## Notes

### Competing Interest Statement

The authors have declared no competing interest.

