## Supplementary material for "Bed and breakfast in the bush: Selection of resting sites and kill sites by leopards (Panthera pardus) on Namibian farmland": Online Resource Material

**Online Resource Material 1:** Leopards used in the study and detailed covariate explanations

**Table S1**: Identification code and sex of captured and collared leopards, time period of data collection, field sampling and types of collars. ● symbol in columns “Resting sites”, “Kill sites” and “Random locations” represents leopards where we sampled data in the field within their home ranges.

| Leopard ID | Sex | Data collection | Resting sites | | Kill sites | Random locations | | | Remote-download GPS collar | Satellite GPS collar | |
| --- | --- | --- | --- | --- | --- | --- | --- | --- | --- | --- | --- |
| L01 | F | 20.5.2014-27.4.2015 |  | ● | | | ● | ● | | |  |
| L02 | M | 21.6.2012-19.2.2014 |  | ● | | | ● | ● | | |  |
| L03 | F | 15.5.2013-27.10.2014 |  | ● | | | ● | ● | | |  |
| L04 | F | 28.5.2013-15.12.2014 |  | ● | | | ● | ● | | |  |
| L05 | M | 25.8.2013-17.5.2014 |  | ● | | | ● | ● | | |  |
| L06 | F | 23.10.2013-1.3.2015 |  | ● | | | ● | ● | | |  |
| L07 | M | 28.10.2013-10.7.2014 |  | ● | | |  | ● | | |  |
| L08 | F | 28.11.2013-17.2.2015 |  | ● | | | ● | ● | | |  |
| L09 | F | 9.12.2013-17.10.2014 |  | ● | | | ● | ● | | |  |
| L10 | M | 9.4.2014-5.12.2015 |  | ● | | |  | ● | | |  |
| L11 | F | 17.4.2014-20.5.2015 |  | ● | | | ● | ● | | |  |
| L12 | M | 3.7.2014-27.7.2016 |  | ● | | |  | ● | | |  |
| L13 | F | 13.7.2014-4.1.2017 |  | ● | | |  | ● | | |  |
| L14 | F | 18.7.2014-5.1.2017 |  | ● | | |  | ● | | |  |
| L15 | F | 26.8.2014-25.10.2014 |  | ● | | | ● | ● | | |  |
| L16 | F | 5.10.2014-5.1.2017 |  | ● | | | ● | ● | | |  |
| L17 | M | 4.12.2014-14.1.2016 |  | ● | | | ● | ● | | |  |
| L18 | M | 6.1.2015-3.3.2015 |  | ● | | |  | ● | | |  |
| L19 | F | 4.6.2015-18.5.2016 |  | ● | | |  | ● | | |  |
| L20 | F | 18.9.2015-8.1.2017 |  | ● | | |  | ● | | |  |
| L21 | F | 2.10.2015-27.7.2016 |  | ● | | |  |  | | | ● |
| L22 | M | 9.12.2015-1.1.2016 |  | ● | | |  |  | | | ● |
| L23 | M | 1.1.2016-9.12.2016 |  | ● | | |  |  | | | ● |
| L24 | F | 2.5.2021-8.1.2022 | ● | ● | | | ● |  | | | ● |
| L25 | F | 14.5.2021-15.4.2022 | ● | ● | | | ● |  | | | ● |
| L26 | M | 1.6.2021-3.9.2022 | ● | ● | | | ● | ● | | |  |
| L27 | F | 20.8.2021-18.6.2022 | ● |  | | | ● | ● | | |  |
| L28 | M | 8.9.2021-28.4.2022 | ● | ● | | | ● |  | | | ● |

**Table S2:** Covariates and data types used in the analyses for leopard resting sites and kill sites with their descriptions. Covariates are categorized into topography, visibility, vegetation and human infrastructure, separated by horizontal line in this order.

| Covariate | Data type | Categories / range | Description |
| --- | --- | --- | --- |
| Relief type | Categorical | Flat | General area (500m in each side from the site) described as flat – no major hills or mountains |
|  |  | Hillside | Hillsides without steep mountains |
|  |  | Mountain | Mountainous areas with steep slopes |
| Special relief features | Categorical | Hillside | Presence of hillside within 30m of the site |
|  |  | Hilltop | Presence of a hilltop within 30m of the location |
|  |  | Ridge | Presence of a ridge within 30m of the location |
|  |  | Small cliff | Presence of a small cliff (maximum height 2m) within 30m of the location |
|  |  | Large cliff | Presence of a large cliff (minimal height 4m) within 30m of the location |
|  |  | Cave | Presence of a cave within 30m of the location |
|  |  | None | No special relief features present within 30m of the location |
| Terrain ruggedness | Categorical | Flat | <20 % within a 20m radius from the site covered with large rocks |
|  |  | Medium rugged | 20-50% within a 20m radius from the site covered with large rocks |
|  |  | Very rugged | >50 % within a 20m radius from the site covered with large rocks |
| Orientation | Categorical | South | Slope facing south |
|  |  | Southeast | Slope facing southeast |
|  |  | Southwest | Slope facing southwest |
|  |  | East | Slope facing east |
|  |  | West | Slope facing west |
|  |  | North | Slope facing north |
|  |  | Northeast | Slope facing northeast |
|  |  | Northwest | Slope facing northwest |
|  |  | Flat | No slope |
| Slope | Continuous | 0 – 89.99 | Inclination on the location in degrees |
| Concealment | Continuous | 0 – 100 | The average concealment calculated by averaging concealments in N, S, E, W; in steps |
| Long-distance view | Continuous | 0 - 365° | Scope of view from the location until 1000m from the site, estimated as a radius of unobstructed view from the site in degrees |
| Long distance viewpoint | Categorical | Elevated | View from site elevated and looking down on the surrounding |
|  |  | Flat | View from site flat |
|  |  | Looking up from below | View from site is not elevated and is looking up from below |
|  |  | None | View obstructed |
| Habitat openness | Categorical | Open | 0-25% of vegetation cover with bushes and trees within 30m radius from the location |
|  |  | Semi open | 26-50% of vegetation cover with bushes and trees within 30 m radius from the location |
|  |  | Dense | 51-100 % of vegetation cover with bushes and trees within 30 m radius from the location |
| Canopy cover | Categorical | Open | 0 % vegetation cover of all vegetation directly above the site |
|  |  | Little open | 1-25% % vegetation cover of all vegetation directly above the site |
|  |  | Semi open | 26-50% % vegetation cover of all vegetation directly above the site |
|  |  | Dense | 51 -100 % vegetation cover of all vegetation directly above the site |
| Shade | Binary | 0/1 | Availability of midday shade of the size of least a leopard body size provided by large tree, bush or cliff in the radius of 30 m from the location |
| Distance to road | Continuous |  | Shortest distance from location to nearest main road in meters |
| Distance to waterhole | Continuous |  | Shortest distance from location to nearest waterhole or dammed water body |
| Distance to house | Continuous |  | Shortest distance from location to nearest occupied farmhouse or house on the outskirts of settlement |

**Online resource material 2:** Statistical analyses for kill sites

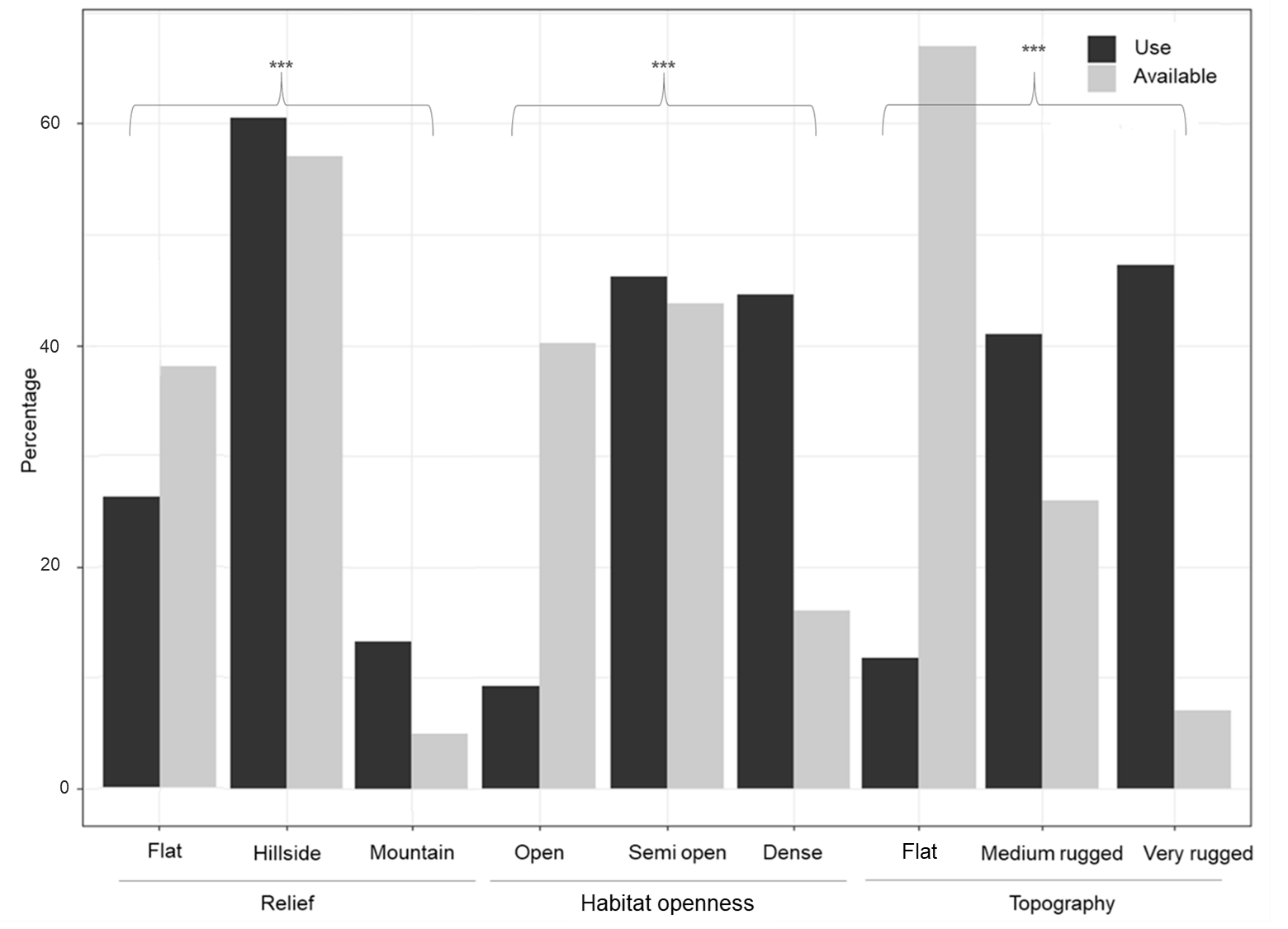

**Fig S1:** Percentage of 195 leopard kill sites (“use”: black) and 195 random locations (“availability”: grey) from 16 leopards for the three covariates relief, habitat openness and terrain ruggedness (each with three categories). All categories were surveyed in the field. *** represents a significant (p < 0.002) difference between “use” and “available”.

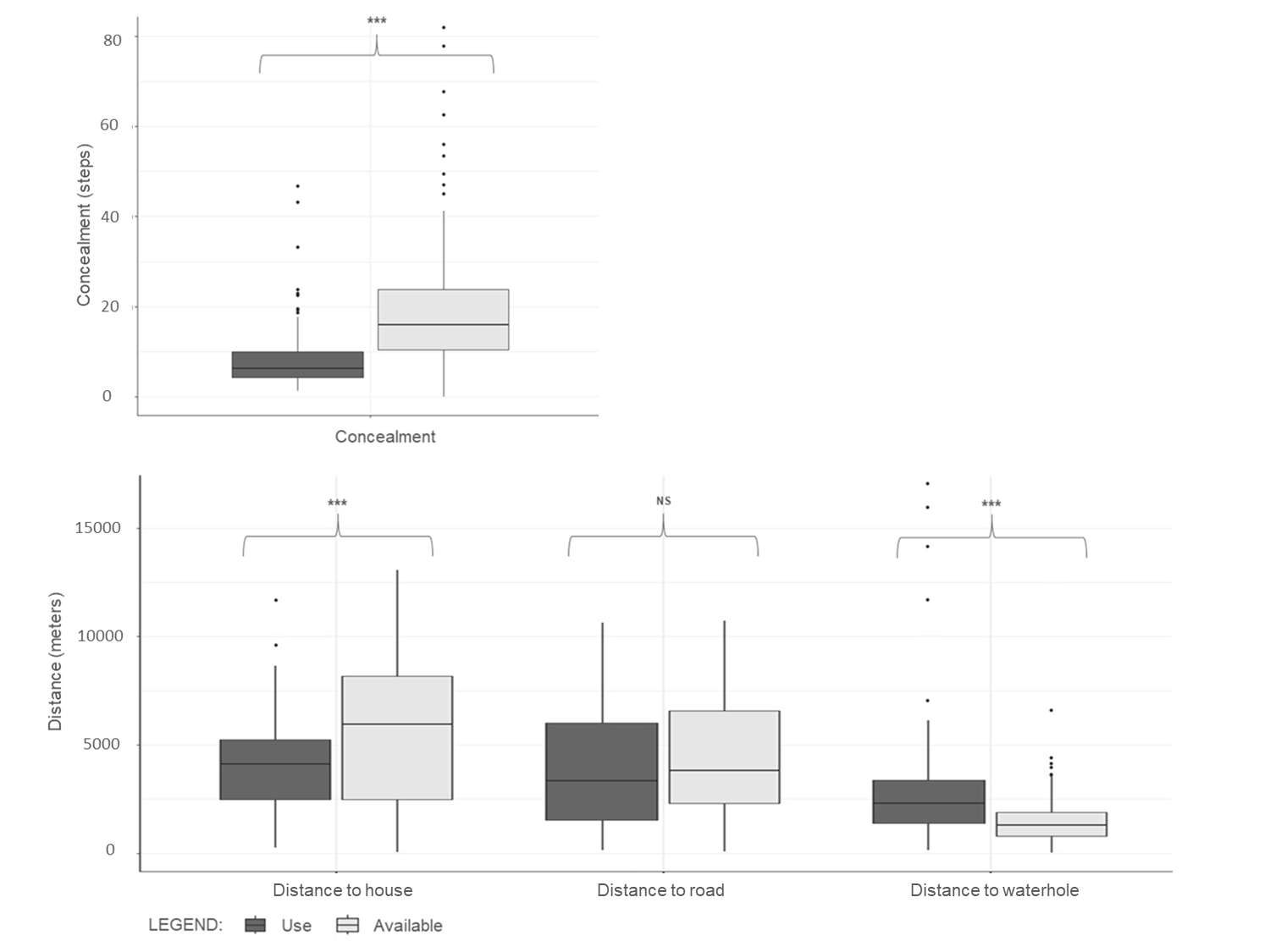

**Fig S2:** Boxplots for continuous covariates of 195 leopard kill sites (“use”: black) and 195 random locations (“available”: grey) from 16 leopards. All categories were surveyed in the field. *** represents a significant (p < 0.001) differences between “use” and “available”.
